# Corpus-wide causality: Algorithm design & application for aggregating gene-disease causal evidence

**DOI:** 10.64898/2026.05.08.723796

**Authors:** Nency Bansal, Adwait P. Parsodkar, Ayush Pathak, Manikandan Narayanan

## Abstract

Identifying causal relationships and distinguishing them from associations is a central scientific endeavor with many applications; knowing causal links between genes and diseases, for instance, can focus drug discovery on curing diseases beyond just symptom management. Despite several studies on automatically extracting relations between entities from large biomedical literature corpora like PubMed, only a few studies extract causal relations from abstracts and even fewer summarize corpus-level evidence for causal links. Recently, Large Language Models (LLMs) have been increasingly deployed to summarize biomedical information and extract relations; however, there is a distinct lack of explicit benchmarking comparing these generalized LLM-based methods against specialized, domain-aware frameworks for corpus-wide causal inference. In this work, we develop a method to infer Corpus-Wide Causal Score (CWCS) of a gene-disease (G-D) pair by integrating two pieces of evidence: (i) network-based causal signals in a prior gene regulatory network, quantified as a CWCS-Net score using an existing multilayer network centrality algorithm; and (ii) corpus-wide literature evidence, quantified as a CWCS-TD (TD for Truth Discovery) score using a newly-developed TD algorithm. Our CWCS-TD (scoring) algorithm jointly and iteratively estimates causal scores for multiple G-D pairs while modeling the reliability of PubMed abstracts co-mentioning them; and represents an advance in the field of TD algorithms due to its incorporation of bibliometric features of publications to address the challenge of sparsity of abstracts that assert a G-D causal relation. Using OMIM as an external expert-curated reference to evaluate classifications of G-D pairs as causal or not, our CWCS method achieved a causal class F1 score of 0.600 across ten diseases, outperforming both LLMs, GPT-4o and MMed-Llama 3 (this performance trend also persists when using area under the precision-recall curve as the evaluation metric). Both LLMs exhibit high recall accompanied by comparatively low precision, resulting in lower causal class F1 scores (0.505 for GPT-4o and 0.522 for MMed-Llama 3) due to large number of false positive predictions. Taken together, these evaluations and other ablation studies show the promise of our carefully designed algorithm in collating and integrating evidence of biomedical causal relations from both network- and literature-based sources, thereby supporting its broader applicability.

## 1. Introduction

Understanding whether a gene causally influences a disease (as opposed to being merely correlated with or responding to the disease) is crucial for targeted drug discovery, personalized medicine, and prioritizing genes for experimental validation. With many published experiments that establish causal genes for various diseases, it is important to collate such causal relations and make them publicly available. This necessitates the development of scalable approaches for comprehensive corpus-wide causal relation extraction, particularly in light of the extensive biomedical literature comprising over 36 million abstracts (Alvarez-Ponce et al., 2026).

While numerous studies have extracted gene-disease (G-D) associations from literature (Bravo et al., 2015; Grissa et al., 2022; Luo et al., 2022) (see also other approaches cited in a survey article (Huang et al., 2024)), only a few have extracted causal relations from individual documents (Gujarathi et al., 2022; Bansal et al., 2024; Lai et al., 2025), and no study (to the best of our knowledge) has automated causal relation extraction of gene-disease pairs from the entire corpus. Simple aggregation approaches such as majority voting often degrade in performance when dealing with more challenging questions (e.g., whether a G-D pair is causal or not) (Aydin et al., 2014), motivating methods that estimate source (e.g., PubMed abstract) reliability rather than relying only on vote counts (Li et al., 2016). Truth Discovery (TD) is one such class of methods that has been widely applied in healthcare (Mukherjee et al., 2014), crowdsourced answering (Li et al., 2020; Ma et al., 2015; Aydin et al., 2014), and information extraction (Li et al., 2014a). Earlier TD methods relied on fixed reputations of sources (Li et al., 2014b), while later techniques used weighted voting and iterative updates (Pasternack and Roth, 2010) as not all papers are equally reliable. However, these approaches do not address source sparsity (Wang et al., 2025), an issue stemming from few papers mentioning the causality of a given G-D pair (due to the significant effort required to establish biomedical causal relations and/or authors’ overcaution in asserting causality).

Furthermore, evidence derived solely from textual literature is often insufficient to establish causal relationships between G-D pairs, as biomedical text frequently contains uncertainty and implicit claims (Yu et al., 2025; Withers et al., 2025). To address this, incorporating prior biological knowledge such as gene-gene interaction (or gene regulatory) networks can provide complementary signals that strengthen G-D relation extraction (Vanunu et al., 2010; Kim et al., 2022; Ata et al., 2021; Deng et al., 2026). To summarize, while literature-driven methods capture aggregated empirical evidence across studies, they often lack mechanistic context; conversely, network-based approaches model underlying biological interactions, but do not fully leverage the breadth of published evidence. Despite their complementary strengths, a unified framework that systematically integrates these two sources of causal evidence remains underexplored.

To address this research gap, we propose a causal scoring framework (Figure 1) with these contributions:

**Fig. 1.**
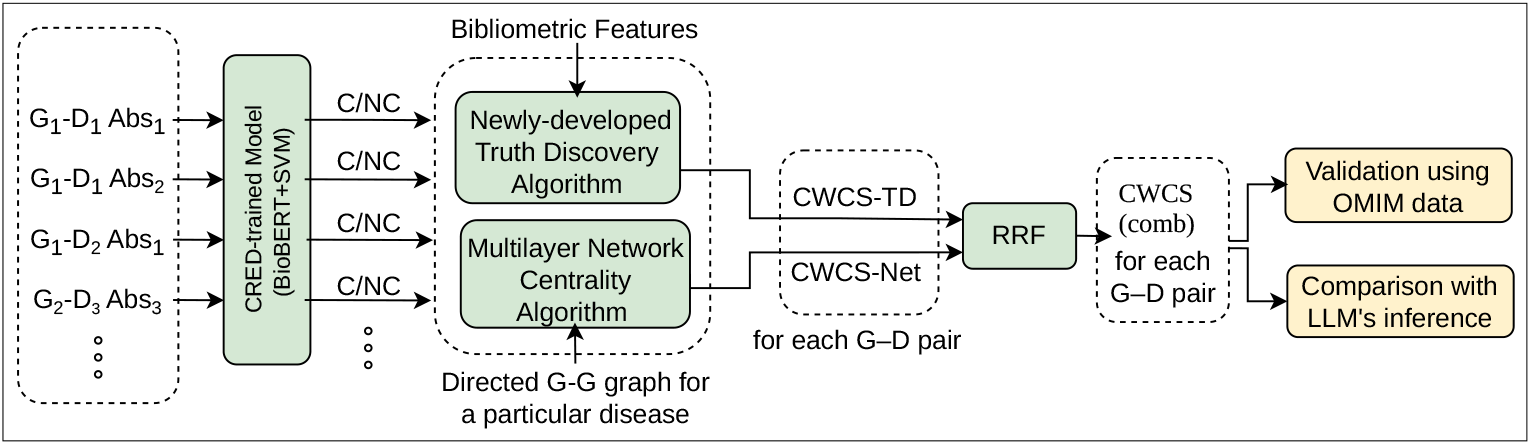
Overview of our work. Note that the same abstract can contain multiple G-D pairs (e.g., Abstract 1) and a G-D pair can be mentioned in different abstracts (e.g., *G*_1_-*D*_1_). C/NC represents the causal/non-causal prediction from CRED-trained model, where CRED denotes Causal Relation Extraction Dataset. RRF refers to Reciprocal Rank Fusion. CWCS (comb) score of a G-D pair quantifies the evidence for G causing D by combining network-based evidence (CWCS-Net) and literature-based evidence (CWCS-TD) scores. Abs denotes Abstract. G-G denotes a gene-gene interaction.

1. We propose an algorithm to compute Corpus-Wide Causal Score (CWCS) to quantify causal G–D relationships by integrating corpus-level literature evidence with network-driven causal influence. To model network effects, we construct a gene–gene (G-G) interaction graph and propagate gene-level information to diseases, resulting in network-based causal scores (CWCS-Net). To capture corpus-wide evidence, we extend the Truth Discovery framework to handle sparse biomedical data, yielding literature-based causal scores (CWCS-TD). These signals are then combined to compute a final corpus-wide causal score (CWCS(comb)) for each G–D pair.
2. We propose CWCS-TD (scoring) algorithm that jointly and iteratively estimates causal scores for multiple G-D pairs while modeling the reliability of PubMed abstracts co-mentioning them. It represents an advance in the field of TD algorithms due to its incorporation of prior information (bibliometric features) and differentiated rewards to respectively address the challenges of sparsity of sources and non-conflicting answers provided by them.
3. We validate the causal scores computed by our framework (CWCS(comb)) using OMIM (Online Mendelian Inheritance in Man) (Amberger et al., 2015) as a reference for ten diseases, demonstrating performance superior to LLM-based baselines.
4. We apply our framework to all the PubMed abstracts for two neurodegenerative diseases – Alzheimer’s disease (AD) and Parkinson’s disease (PD) – to identify disease-causing genes beyond those present in the reference database.

Given the promising results, our proposed algorithm (available at https://github.com/BIRDSgroup/CWCS) can be widely applied on the PubMed corpus of abstracts to find causal genes of other diseases as well.

## 2. Methods

We propose a method to quantify the causality of G-D relationships by integrating network-driven causal influence with corpus-wide literature evidence. The algorithm to compute this CWCS score operates by first constructing a disease-specific directed network that captures both G–D and G–G regulatory relationships in different layers. Causal influence scores are then computed using a PageRank-like multilayer network centrality (also called MultiCens) algorithm, and subsequently aggregated with literature-derived causal evidence obtained via a truth discovery mechanism. CWCS-Net refers to the network-based framework built upon MultiCens, or the corpus-wide causal scores derived from it (the term’s usage should be clear from the context). Similarly, CWCS-TD denotes the newly-developed Corpus-Wide Causal Scoring Truth Discovery algorithm developed in this study, or the corresponding corpus-wide causal scores inferred under this formulation.

### 2.1. Causal score calculation using CWCS-Net algorithm

We apply the centrality measures of MultiCens framework (Kumar et al., 2023), referred to as the CWCS-Net algorithm, to quantify the network-based causal influence of genes on the disease. We apply MultiCens on each disease separately to compute the causal effect of the gene directly on the disease (locally) and through other genes (globally). To compute this effect/score for a given disease, we need to first assemble a list of G→D and G→G relationships, which are then used to construct an appropriate multilayer network on which MultiCens can be applied.

#### 2.1.1. Assembling directed edges involving genes and a disease

##### Source of Gene→Disease (G→D) edges

Directed edges from genes to the disease node are constructed for all genes present in the OMIM page for the corresponding disease. Edge weights represent the strength of causal association and are computed as the mean probability predicted by a CRED-trained BioBERT+SVM model across all abstracts that mention the given G–D pair. This probabilistic weighting captures literature-derived evidence of causality while smoothing noise from individual predictions. As a brief background, note that CRED is a manually-annotated benchmark dataset of disease-causing genes identified in PubMed abstracts, and moves beyond simple co-occurrence or association of G-D pairs. The creation of CRED involved a multi-stage annotation process, wherein experts examined biomedical literature to annotate G-D pairs to be causal/non-causal in the given abstract, based on linguistic cues and other pieces of evidence in the abstract. CRED enjoys a high inter-annotator agreement (Cohen’s Kappa = 0.89). For the predictive component of this study, we employed a classification model trained on CRED (called BioBERT+SVM model) that was found to outperform other CRED-trained models including LLMs. For more details regarding CRED and the model trained on CRED, please refer to our earlier study (Bansal et al., 2024).

##### Source of Gene→Gene (G→G) edges

Pairwise directed regulatory interactions among genes (e.g., transcription factor to target gene interactions) are obtained from OmniPath, a comprehensive meta-database of experimentally validated and literature-curated interactions of different types (such as signaling, regulatory, and protein-protein interactions) integrated from over 100 molecular biology resources (Türei et al., 2026). OmniPath reports the number of different databases supporting a specific interaction, which allowed us to filter for only “high-confidence” links; and also provides the directionality of relations among entities, which allowed us to focus on only directed (causal/regulatory) interactions. Each directed G–G edge weight is defined as the number of independent sources supporting the regulation, normalized by the maximum number of supporting sources observed across the OmniPath dataset (OmniPath, 2015). Number of G-D and G-G edges in the directed graph of each disease are shown in Table S2 in the Appendix.

#### 2.1.2. CWCS-Net Algorithm

For a given disease, we construct its multilayer network, and represent this directed network as a supra-adjacency matrix constituted from two matrices (A for capturing within-layer connections alone, and C for across-layer connections); and then use these two matrices to define local and global centrality measures. Specifically, we adapt the MultiCens framework (designed to capture node importance in multilayer networks with inter-layer coupling (Kumar et al., 2023)) to a G-D causality setting by interpreting the multilayer structure as a two-layer network: a) G–D layer capturing directed relation between genes and the disease, and b) G–G layer capturing regulatory interactions among genes. Supra-adjacency matrix capturing both intra- and inter-layer connections involving these two layers is given by:

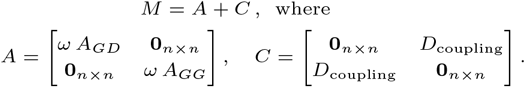

Here,

- *A*_*GD*_ and *A*_*GG*_ respectively represent the row-normalized adjacency matrices of the directed network of G→D and of G→G edges assembled in the previous section. The dimension of *A*_*GD*_ and *A*_*GG*_ is *n* x *n* each, where *n* is the number of nodes in each layer of the multilayer network (*n* − 1 genes and 1 disease node).
- *ω* ∈ [0, 1] refers to a coupling constant (set to *ω* = 0.5 in our analyses to give equal weight to the G-D vs. G-G layers).
- *D*_coupling_ refers to an *n* × *n* diagonal matrix capturing the inter-layer coupling between the G–D and G–G layers, and is constructed using the coupling weight of node *i* as:

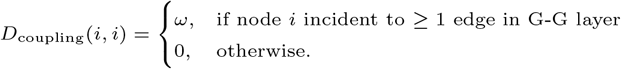

Note that the set of G→D and of G→G edges assembled in the previous section “as is” may result in unnormalized adjacency matrices, which are then row-normalized (i.e., each row’s sum, denoted rowsum, converted to 1) as follows. Each row of the unnormalized A_*GD*_ has been normalized by adding a self-loop of weight 1-rowsum to the diagonal; and of unnormalized A_*GG*_ has been normalized by adding a self-loop of weight 1-rowsum to the diagonal if rowsum < 1, or dividing each edge weight by rowsum otherwise).

The supra-adjacency matrix *M* is finally row-normalized, i.e., each entry in the matrix divided by the corresponding row’s sum, to obtain 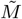 whose rows sum to 1. After decomposing,

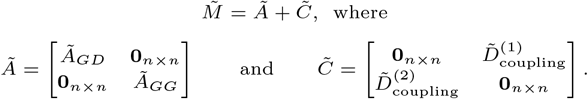

To capture the influence of a gene locally and globally, we respectively compute *local* centrality which captures the influence of a node within its own layer, and *global* centrality that reflects the node’s influence across all layers in the multilayer network. The local centrality of the *n* nodes in layer *i* (denoted by the length-*n* vector 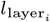 where layer_*i*_ ∈ {*GG, GD*}) is defined recursively and computed iteratively (Kumar et al., 2023) as:

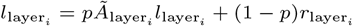

Here, (1−*p*) := 0.25 refers to the restart probability, and 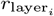is a length-n restart vector that is set to 1 for the disease node and 0 for all other nodes. A uniform restart vector was used in the original MultiCens formulation, however a disease-specific restart vector is used here to bias a random walker on this multilayer network towards the disease of interest.

The global centrality vector (length-2*n* vector *g* := [*g*_*GD*_; *g*_*GG*_]) can then be computed using the length-2n local centrality vector *l* := [*l*_*GD*_; *l*_*GG*_] computed above, and the length-2*n* restart vector 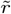 defined as 1 for only the disease node in the G-D layer and 0 for all other nodes as:

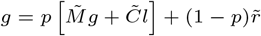

The CWCS-Net score for a gene *j* is then simply taken as the aggregate of its global centrality across both the layers, in order to capture the directed connectivity of the gene in both layers to the disease node in the G-D layer, i.e.,

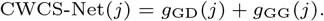

### 2.2. Causal score calculation using CWCS-TD algorithm

In this section, we begin with the principles of traditional Truth Discovery, where numerous sources provide with conflicting evidences that need to be aggregated, and extend the algorithm to our setting (referred as CWCS-TD) to address these key challenges: (i) only a few sources (PubMed abstracts) co-mention a given G–D pair and answer the question of whether the pair is causal, and (ii) a source’s response to the causal question is either “yes” or “maybe” rather than a definitive and conflicting “no”.

#### 2.2.1. Traditional truth discovery approach

It is important to note that reliability annotations for sources are not provided *a priori*; instead, the algorithm is tasked with inferring them solely based on the sources’ responses. Truth Discovery approaches this challenge through a circular definition (Parsodkar et al., 2023):

> *An answer is deemed trustworthy if it is supported by several reliable sources, while a source is considered reliable if it provides trustworthy answers to several questions*.

This formulation encapsulates the mutual dependency between the trustworthiness of answers and the reliability of the sources that provide them.

The resolution of this circularity has been approached through a variety of strategies, ranging from optimizing a weighted distance function (Li et al., 2014b) to methods involving Probabilistic Graphical Models (Ma et al., 2015). To realize our contribution in this work, we build upon an existing optimization-based framework proposed by (Li et al., 2014b), shown below:

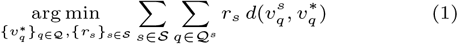

Here, *d*(·, ·) is a distance function and, *S* and *Q* denote the set of sources and the set of questions, respectively. We use *S*^*q*^ to denote the sources that answer *q* ∈ *Q* and *Q*^*s*^ to denote questions answered by source *s* ∈ *S*. For all notations refer to Table S1 in the Appendix. The optimization function in Equation 1 involves estimation of two sets of parameters: the reliability of sources 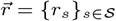, and the trustworthy answers to questions, also referred to as identified truths, 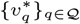. To prevent trivial solutions—such as assigning r_*s*_ = −∞ for all *s* ∈ *S*—we constrain the reliability vector 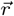 to lie within a feasible set Δ. We refer interested readers to refer (Li et al., 2014b; Parsodkar et al., 2022) for instances of different feasible regions used in related work.

#### 2.2.2. Extension of truth discovery algorithm

In this section, we develop a CWCS-TD algorithm by modifying the objective function above to handle sparsity using prior information, and to handle non-conflicting (yes/maybe rather than yes/no) answers from sources using differentiated rewards. While Truth Discovery has been employed in a variety of settings for assessing source reliability, existing approaches typically begin with no prior indicators of source quality. We posit that in certain settings, such as our sparse setting, it is both feasible and beneficial to incorporate prior knowledge about sources, which can guide the TD process toward more meaningful and accurate estimates of source reliability.

For each source *s* ∈ *S*, we represent its associated information *x*^*s*^ as a set of feature–value pairs, i.e., 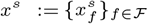. In our use case, where PubMed articles (research papers) are sources, we selected three features to estimate paper reliability: i) h-index of the journal in which the paper is published, ii) years since published (hereby referred to as publication age), and iii) citation count. A journal’s h-index indicates that it has published *h* papers that have each been cited at least *h* times, indicating the volume of impactful, frequently cited work. Citation count serves as a proxy for the scientific influence and acceptance of a study within the research community. Publication age is incorporated to account for the fact that older studies have been available long enough for the scientific community to critically examine their findings. Together, these features provide an interpretable measure of reliability by integrating venue quality, sustained influence, and time-based stability. All feature values are normalized using min–max scaling to ensure they fall within the interval [0, 1]. Moreover, we assume that the features exhibit a *more-is-better* property: increasing a feature value either enhances or does not affect the reliability of the source, but never diminishes it.

Given access to feature values for all sources, we model the reliability of a source as a linear combination of its feature values, weighted by *β* = {*β*_*f*_}_*f*∈ℱ_, which captures the relative importance of each feature:

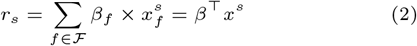

Furthermore, we assign differentiated rewards to distinct answers. This design choice is motivated by scenarios where candidate answers are *non-orthogonal* — that is, when a source supporting one answer does not necessarily oppose the others. Consider the label set `ℒ = {yes, maybe`} where a source providing the answer `maybe` partially supports `yes`. To discourage sources from consistently opting for the *safer* response (`maybe`), we introduce higher rewards for providing a decisive answer (`yes`, in this case) when it aligns with the identified truth. To formalize this, we introduce a hyperparameter λ ∈ (0, 1) that discounts weaker answers.

By substituting this formulation into Equation 1 and reformulating the objective in terms of maximizing similarity rather than minimizing distance, we obtain a new optimization function over the parameters β = {β_*f*_}_*f*∈ℱ_ and 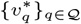, expressed as:

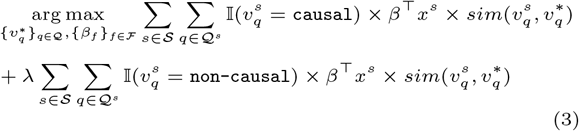

where 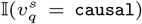 returns 1 if source *s* claims question *q* to be causal, and 0 otherwise. Additionally, we impose the constraint on *β* rather than on 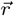, such that 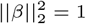.

#### 2.2.3. Optimization Strategy

In this section, we focus on the optimization strategy for the proposed objective function within a binary classification setting. We model the claims made by sources as:

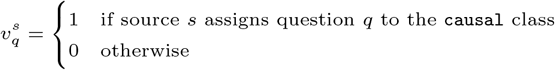

This strategy enables us to interpret the aggregated answer for a question *q* as a measure of confidence in assigning *q* to the `causal` class. Specifically, 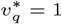 indicates high confidence in *q* belonging to the `causal` class, while 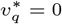 suggests otherwise. Based on this interpretation, we define the similarity function as follows:

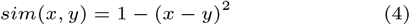

The minimization of the objective function involves an Expectation-Maximization style optimization that involves the optimization of one set of parameters in light of the knowledge of the other. In our case, the *T*-step (responsible for estimating the values 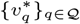, see Section CWCS-TD Algorithm in Appendix) computes the set of trustworthy answers conditioned on the knowledge of the source reliability scores as follows. Let 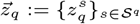. Also, let

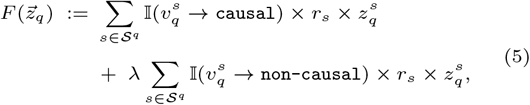

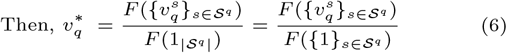

The *R*-step (responsible for estimating the reliability scores of sources, see Section CWCS-TD Algorithm in the Appendix), on the other hand, estimates the reliability of sources contingent on the identified truths. Since *β* serves as a precursor for estimating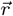, we employ the following equations to determine the *β* values:

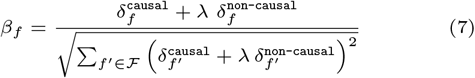

where, 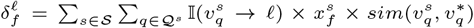 and *ℓ* ∈ {`causal, non-causal`}. Given the {*β*_*f*_}_*f*_ values, the {r_*s*_}_*s*_ values can be computed using Equation 2. The derivation of these equations is elaborated in Section CWCS-TD Algorithm in the Appendix. Algorithm 1 in the Appendix shows our complete CWCS-TD algorithm.

#### 2.2.4. Learning and application of β

The CWCS-TD algorithm described above (Algorithm 1 in the Appendix) was first applied to the CRED dataset (containing PubMed abstracts with ground-truth labels of G-D causal/non-causal pairs) to estimate the reliability feature weights *β*. The learnt *β* values were: 0.231, 0.644 and 0.729 for citation count, journal h-index, and publication age respectively. These learnt weights are used to compute the reliability score of each paper (using Equation 2), and subsequently the corpus-wide causality score of each G-D pair considered in the OMIM validation corpus or other downstream applications (using Equation 6).

### 2.3. Aggregation of corpus-wide causal scores

We aggregate the causal scores obtained from CWCS-Net and CWCS-TD components using the Reciprocal Rank Fusion (RRF) technique (Cormack et al., 2009). RRF is a rank-based aggregation method that combines evidence from multiple scoring functions by prioritizing entities that are consistently ranked highly across methods, while remaining robust to score-scale differences. It has been validated to outperform other fusion techniques like Borda count in settings where the underlying scoring mechanisms are fundamentally different (Benham and Culpepper, 2017).

For each gene corresponding to a disease, ranks are independently computed based on the CWCS-Net derived causal scores and the CWCS-TD derived causal scores. The final aggregated causal score of each gene *j* is computed as the sum of the reciprocal ranks from the two methods, using the following formula, where k was set to default value of 60.

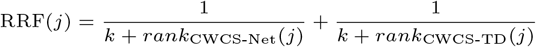

The computed RRF scores are scaled using quantile transformer (from sklearn library), that tends to spread the most frequent values and reduce the impact of outliers, without impacting the ranking. The final RRF score assigns higher importance to genes that are ranked highly by both network-based and literature-based inference, while still allowing genes strongly supported by either component to contribute meaningfully to the final ranking.

### 2.4. Validation using OMIM

To evaluate corpus-wide causal scores of G-D pairs, we use expert-curated G-D pairs from the OMIM database (Amberger et al., 2015) as reference ground truth. OMIM can be regarded as a “gold standard” due to its rigorous expert-driven curation process – unlike automated databases, OMIM entries are manually synthesized from primary biomedical literature, wherein curators critically evaluate clinical evidence, phenotypic characteristics, and inheritance patterns before establishing gene–phenotype associations. We selected ten diseases with the highest number of OMIM-listed causal genes, as summarized in Table 1.

**Table 1.**
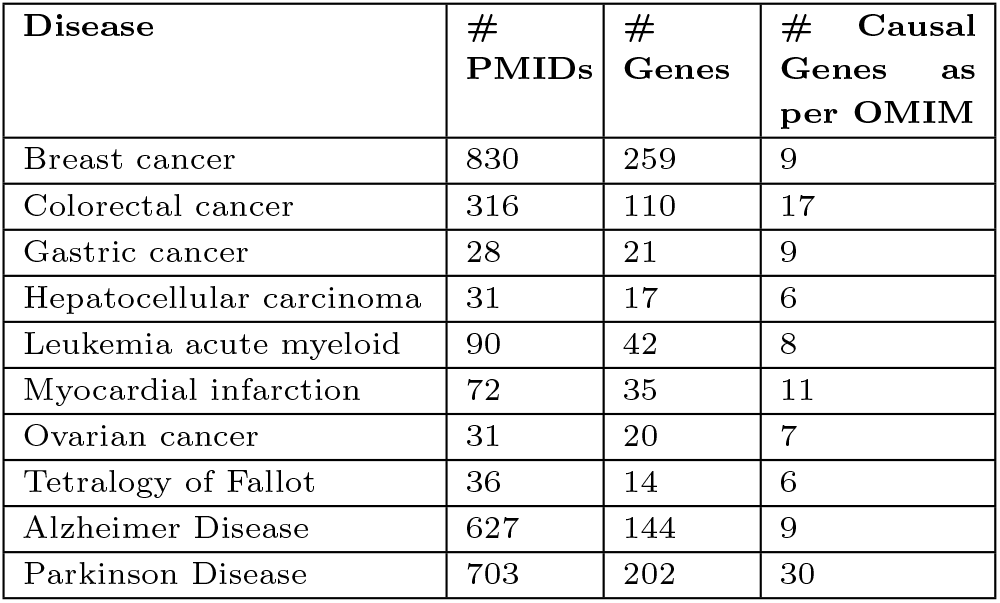
Statistics of OMIM validation corpus. Please see text for details. # stands for “Number of”.

| Disease | # PMIDs | # Genes | # Causal Genes as per OMIM |
| --- | --- | --- | --- |
| Breast cancer | 830 | 259 | 9 |
| Colorectal cancer | 316 | 110 | 17 |
| Gastric cancer | 28 | 21 | 9 |
| Hepatocellular carcinoma | 31 | 17 | 6 |
| Leukemia acute myeloid | 90 | 42 | 8 |
| Myocardial infarction | 72 | 35 | 11 |
| Ovarian cancer | 31 | 20 | 7 |
| Tetralogy of Fallot | 36 | 14 | 6 |
| Alzheimer Disease | 627 | 144 | 9 |
| Parkinson Disease | 703 | 202 | 30 |

#### 2.4.1. OMIM validation corpus and task

We validated corpus-wide scores using a OMIM validation corpus assembled for each of the ten diseases listed in Table 1. In detail, for any given disease of interest, we extracted rows/entries from the `genemap2.txt` file provided by OMIM, and retained only rows in which the phenotype corresponded to the disease and was associated with at least one gene. Note that each of these retained rows/entries is associated with a unique MIM number, a 6-digit identifier corresponding to a manually curated OMIM record describing a causal G-D relationship supported by experimental or literature evidence. We then collected PubMed identifiers (PMIDs) referenced by all retained MIM numbers associated with the disease via automated web scraping (OMIM, 2015). Abstracts corresponding to these PMIDs were retrieved using the PubTator3 API from the National Center for Biotechnology Information (NCBI), which performs named entity recognition to map gene mentions to Entrez identifiers and disease mentions to MeSH identifiers. The number of PMIDs obtained for the disease and of genes recognized across the corresponding abstracts are reported in Table 1. This overall data is referred to as the OMIM validation corpus in this paper.

For each G-D pair in the above OMIM validation corpus, the validation task is to predict if it’s causal or not based on all abstracts in the corpus co-mentioning this G-D pair. To aid this validation task, we perform causal relation extraction on each relevant abstract as follows: gene and disease mentions in the abstract were masked as `GeneSrc` and `DiseaseTgt` respectively, and the masked abstracts were provided as input to a CRED-trained BioBERT+SVM model (Bansal et al., 2024). The resulting binary predictions (`causal` vs. `non-causal` predicted labels) were provided as inputs to CWCS-TD, and the associated prediction probabilities were provided as inputs to CWCS-Net. Both prediction labels and probabilities were provided as inputs to the LLM-based models.

#### 2.4.2. Evaluation of the CWCS algorithm

For evaluating CWCS-Net algorithm on the OMIM validation task, we apply the algorithm separately on each of the ten diseases of interest using relevant inputs. For evaluating CWCS-TD algorithm on the OMIM validation task, we run it separately on each G-D pair (as described in Section 2.2.4). To do so, for the abstracts in the OMIM validation corpus co-mentioning the G-D pair, the CWCS-TD algorithm requires as input the binary prediction labels from the CRED-trained model mentioned above, as well as the bibliometric features like citation count, publication age, and h-index of the abstracts to support reliability estimation. Citation count and publication age were obtained via the Semantic Scholar API (SemanticScholar, 2025), while h-index values were retrieved from SCImago (SCImago, 2025).

### 2.4.3. Evaluation of LLM-based inference

To assess the capability of LLMs to score the support for causal relationship between genes and diseases in biomedical literature, we evaluated two state-of-the-art models: MMed-Llama 3 (MMed-Llama-3-8B) (Wu et al., 2024) and GPT-4o (OpenAI, 2025). MMed-Llama-3, a medical domain-specific language model, was deployed locally, while GPT-4o inference was performed using OpenAI API services. For evaluating a LLM-based model on the OMIM validation task, we run it separately on each G-D pair using a system prompt that instructed the model to act as a biomedical expert and analyze all relevant abstracts (the same set of abstracts provided as input to CWCS-TD) to determine support for the G-D pair’s causal relationship. The prompt explicitly requested responses in the specific format `<relationship>: <score>`, where `<relationship>` was either `Causal` or `Not causal`, and `<score>` represented a confidence value between 0 and 1. In detail, the user prompt presented the aggregated text from all relevant abstracts for the G-D pair, along with CRED-trained model predictions for the abstracts as mentioned above, and asked the model to determine whether a causal relationship exists between the entities labeled as “GeneSrc” and “DiseaseTgt”. The user prompt also clarified that associations are not necessarily causal relationships and instructed the model to look for explicit mentions of causality (for the exact prompt provided to LLMs, see Section LLM Inference: Prompt to LLM in the Appendix). To ensure the reliability of the extracted predictions, we validated responses to identify malformed outputs that required manual review.

## 3. Results

### 3.1. Inferred relationship between bibliometric features and reliability score using the manually annotated CRED

As seen in Methods, CWCS-TD component of our CWCS framework computes the reliability score of each paper along with the corpus-wide causal score of each G-D pair. Reliability score of each research paper depends on three key bibliometric features: h-index, publication age, and citation count. Figure 2 illustrates the relationship between the reliability score and these bibliometric indicators, based on CRED dataset. The h-index shows a high correlation with reliability scores, implying that journals with high productivity and citation impact are more likely to publish papers with reliable and verifiable findings. Publication age is also highly correlated with the reliability score, whereas the citation count shows only a modest positive correlation with the reliability score. This indicates that while citations may contribute to reliability estimation, they alone are not sufficient predictors. Citation counts are also influenced by external factors such as the field’s size and topical popularity. Hence, all the features together must be used for more accurate assessment of research reliability.

**Fig. 2.**
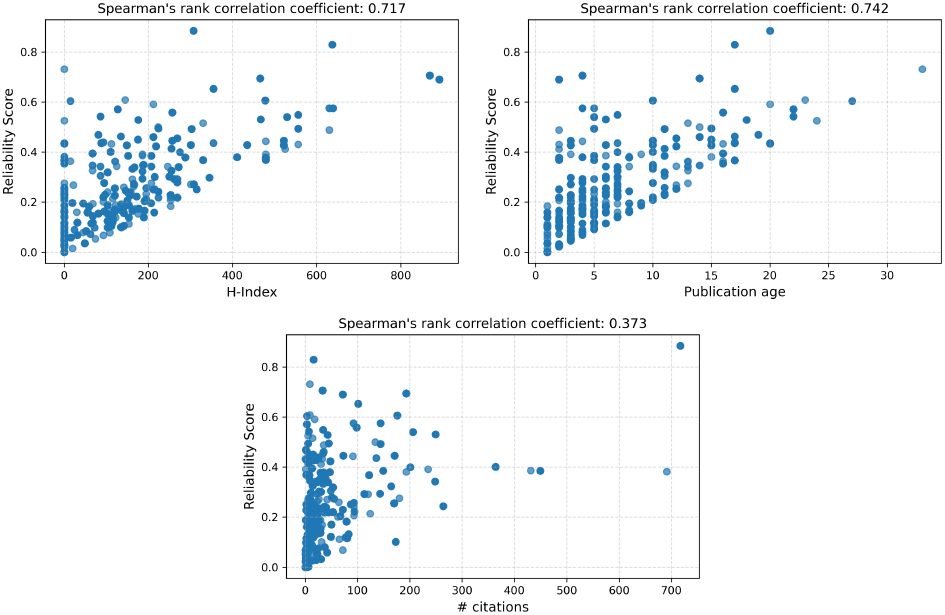
Relationship between each bibliometric feature and the inferred reliability score.

### 3.2. Validation using OMIM

All validation results are reported using the OMIM validation corpus constructed for the ten diseases described in Methods. Our CWCS method, along with other evaluated methods, was applied to the OMIM validation corpus dataset underlying Table 1. But the evaluation metrics were computed only for genes found in more than one publication within the OMIM validation corpus (see Table S3 in the Appendix), to ensure adequate literature coverage for the benchmarked methods aggregating corpus-wide evidence.

#### 3.2.1. Evaluation and comparison of the CWCS algorithm

We evaluated the CWCS framework on the OMIM validation corpus across ten diseases and compared its performance against individual components, baseline classifiers, LLMs, and alternative aggregation strategies (see Table 2). It includes comparison with a ranking approach CausalRatio based on the causal ratio, which is defined for each G-D pair as the ratio of the number of papers in which this pair is predicted as causal (by the CRED-trained model) to the total number of papers mentioning this G-D pair.

**Table 2.** Comprehensive performance comparison including PR-AUC and other classification metrics for all causal gene prediction models. For each method, the reported cutoff is decided based on the maximum F1 score of the causal class. In the Model column, “A+B” refers to combining scores from Model A and B using the RRF method (e.g., CWCS-Net+CWCS-TD is basically CWCS (comb) or CWCS simply).

| Model | Causal class |  |  |  | Macro average |  |  | PR-AUC |
| --- | --- | --- | --- | --- | --- | --- | --- | --- |
|  | cut-off | Precision | Recall | F1 Score | Precision | Recall | F1 Score |  |
| Random Classifier | 0.01 | 0.309 | 1.000 | 0.472 | 0.154 | 0.500 | 0.236 | 0.309 |
| CausalRatio | 0.11 | 0.389 | 0.783 | 0.520 | 0.606 | 0.617 | 0.552 | 0.440 |
| CWCS-TD | 0.85 | 0.389 | 0.783 | 0.520 | 0.606 | 0.617 | 0.552 | 0.420 |
| GPT-4o | 0.27 | 0.345 | **0.940** | 0.505 | 0.614 | 0.572 | 0.418 | 0.535 |
| MMedLlama-3 | 0.62 | 0.408 | 0.723 | 0.522 | 0.610 | 0.628 | 0.582 | 0.440 |
| CWCS-Net | 0.46 | 0.425 | 0.747 | 0.541 | 0.627 | 0.648 | 0.601 | 0.428 |
| CWCS-Net+CausalRatio | 0.39 | 0.394 | 0.783 | 0.524 | 0.610 | 0.623 | 0.559 | 0.472 |
| CWCS-Net+GPT-4o | 0.48 | 0.421 | 0.711 | 0.529 | 0.618 | 0.638 | 0.598 | 0.510 |
| CWCS-Net+MMedLlama-3 | 0.37 | 0.396 | 0.807 | 0.532 | 0.618 | 0.629 | 0.560 | 0.431 |
| CWCS-Net+CWCS-TD | 0.64 | **0.557** | 0.651 | **0.600** | **0.694** | **0.710** | **0.699** | **0.601** |

Among all evaluated methods, CWCS (comb) achieves the best overall performance, with causal class F1 score of 0.600 and PR-AUC of 0.601. Despite CWCS-TD’s and CWCS-Net’s comparatively weaker standalone performances, aggregating both using RRF in CWCS (comb) consistently improves causal gene prioritization – specifically by an absolute PR-AUC gain of ~17% – indicating that corpus-level evidence and network topology provide complementary information. These trends are further reflected in the precision-recall (PR) and Recall@k curves shown in Figures 3 and 4. CWCS-Net+CWCS-TD consistently improves the ranking of known causal genes, particularly among top-ranked candidates, demonstrating that corpus-wide reliability modeling refines network-driven causal inference rather than replacing it.

**Fig. 3.**
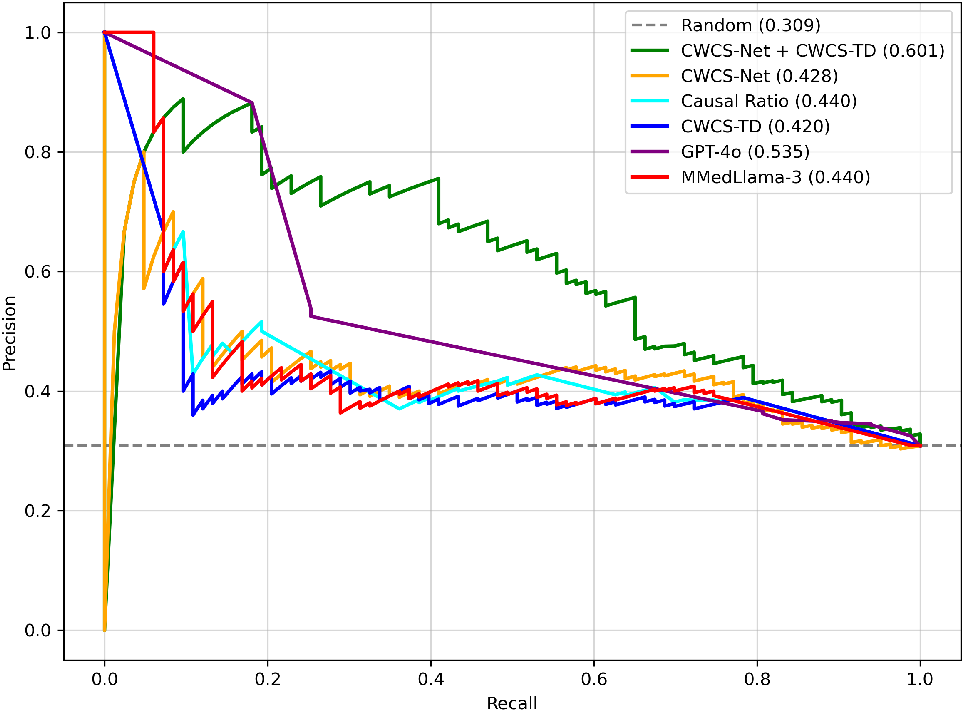
PR curves of different models on OMIM validation corpus. See also Figure S1 in the Appendix for PR curves of other fusion methods.

**Fig. 4.**
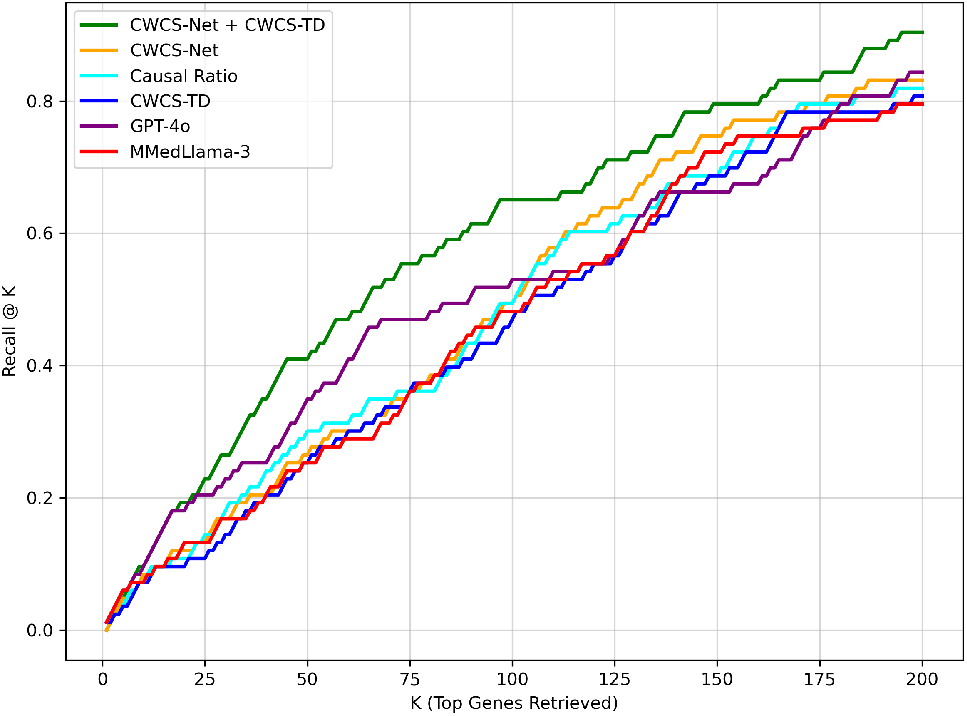
Recall@K curves of different models on OMIM validation corpus. See Figure S2 in the Appendix for similar plots of other fusion methods.

In addition to recovering known causal genes, CWCS identifies candidate genes that are not currently annotated as causal in OMIM. Among the top 30 top-scoring (CWCS (comb)) G-D pairs, 22 were validated as true positives against OMIM. Of the 8 pairs not supported by OMIM, manual inspection showed published causal evidence for *MLLT10* →AML and *STAT5A* →AML (here, AML refers to Acute Myeloid Leukemia). In contrast, no direct studies were identified for the predicted causal links from *ESR1* and *GSK3B* to Ovarian cancer; however, both genes regulate OMIM-listed causal genes for Ovarian cancer such as *AKT1, CDH1*, and *CTNNB1* (suggesting that their high scores arise from strong regulatory evidence captured by the network component; see Table 3). The remaining 4 pairs may be treated as false positives (FPs). Overall, these findings demonstrate the ability of CWCS to integrate complementary evidence sources and surface biologically relevant candidates beyond curated databases.

**Table 3.** Of the 30 G-D pairs with the highest CWCS (comb) scores, the 8 listed here are not captured in OMIM, and could hence be potentially novel causal links (supported by textual evidence (PMID) or regulatory network edges; see Remarks column) or likely false positives (FPs). TD, Net and RRF denote CWCS-TD, CWCS-Net and CWCS (comb) scores respectively.

| Gene | Disease | TD | Net | RRF | Remarks |
| --- | --- | --- | --- | --- | --- |
| *MLLT10*<br>/*AF10* | AML | 0.990 | 0.976 | 0.860 | PMID: 8643484 |
| *STAT5A* | AML | 0.991 | 0.872 | 0.843 | PMID: 16410449 |
| *ESR1* | Ovarian cancer | 0.000 | 0.411 | 0.842 | *ESR*→*AKT1*,<br>*CDH1*, *CTNNB1* |
| *GSK3B* | Ovarian cancer | 0.000 | 0.272 | 0.817 | *GSK3B*→*AKT1*,<br>*CDH1*, *CTNNB1* |
| *KMT2A*/<br>*MLL* | AML | 0.973 | 0.951 | 0.834 | Likely FP |
| *TP53* | AML | 0.996 | 0.755 | 0.829 | Likely FP |
| *AP2B1* | Myocardial infarction | 0.976 | 0.964 | 0.990 | Likely FP |
| *WWOX* | Ovarian cancer | 0.990 | 0.660 | 0.940 | Likely FP |

#### 3.2.2. Evaluation of LLM-based inference

To contextualize the performance of our CWCS framework, we compared it against the performance of two LLMs, MMed-Llama 3 (Wu et al., 2024) and GPT-4o (OpenAI, 2025); and report the results in Table 2, and Figures 3 and 4. Both LLMs exhibit high recall on the OMIM validation corpus, with GPT-4o achieving a recall of 0.94 for the causal class. This behavior indicates strong sensitivity to potential causal mentions in biomedical text. However, the high recall is accompanied by comparatively low precision, resulting in lower causal F1 scores (0.505 for GPT-4o and 0.522 for MMed-Llama 3) and PR-AUC values (0.535 and 0.440, respectively). Analysis of the confusion matrix entries (Table S4 in Appendix) confirms that this performance is driven by a large number of FP predictions when LLMs are applied directly to corpus-level causal inference.

### 3.3. Application of CWCS to all AD/PD PubMed abstracts and its interpretation

To assess the practical utility of our CWCS method, we applied it to the complete set of PubMed abstracts co-mentioning at least one gene and a disease of interest (Alzheimer’s disease (AD) or Parkison’s disease (PD)), using the same parameters/weights as in the validation experiments. This full PubMed corpus, which extends beyond the OMIM validation corpus, included abstracts published up to March 2026; and amounted to 71,586 abstracts covering 10,574 genes for AD, and 51,486 abstracts covering 12,926 genes for PD.

Applying CWCS to the full PubMed corpus results in improved recovery of certain OMIM-listed causal genes for both diseases. In the case of AD, some genes curated as causal in OMIM but weakly supported in the OMIM validation corpus receive higher causal scores when all PubMed abstracts are considered. For example, *ADAM10*, which initially received a CWCS (comb) of 0.327 in the OMIM validation corpus, attains a high CWCS of 0.569 when applied on the full corpus. This increase reflects the incorporation of additional studies that explicitly describe the causal role of *ADAM10* in the pathogenesis of AD and its regulatory interactions.

Similar trends were observed for PD. Genes such as *VPS35* and *PLA2G6*, which were not prioritized by CWCS as causal using the OMIM validation corpus (CWCS (comb) of 0.206 and 0.183, respectively), achieve high CWCS of 0.836 and 0.798 respectively when the complete PubMed corpus is analyzed. These improvements indicate that CWCS effectively aggregates distributed causal evidence that is sparse at the curated-database level but consistent across the broader literature.

We manually verified the top 10 (CWCS) scoring genes for both AD and PD. We found that for AD, 2 out of these 10 are true positives (i.e., also reported in OMIM as causal), 5 can be potentially novel causal candidates based on textual/regulatory evidence (see Table S5 in the Appendix), and the remaining 3 genes may be FPs. For PD, 9 out of the top 10 genes are true positives, and the remaining gene may be a FP (see Table S6 in the Appendix). We also manually verified a subset of abstracts with high estimated reliability scores, and understood the excerpts from such abstracts that contributed positively or negatively to the prediction of OMIM-listed causal genes for AD (see Table S7) and for PD (see Table S8).

Overall, these findings demonstrate that incorporating the complete PubMed corpus enables CWCS to detect both well-established and sparsely studied causal genes. By aggregating signals across the literature while leveraging regulatory network structure, CWCS improves sensitivity to dispersed causal evidence without compromising consistency for well-characterized disease genes.

## 4. Conclusion

Truth Discovery methods provide a principled mechanism for aggregating evidence from multiple sources, but their effectiveness is often limited by data sparsity. This challenge is particularly pronounced in biomedical causal relation extraction, where explicit causal claims are rare, inconsistently reported, and distributed across a small subset of publications. As a result, corpus-level aggregation based solely on textual evidence is often noisy and incomplete, even for well-established gene-disease relationships.

To address these challenges, we propose an algorithm to compute Corpus-Wide Causal Score (CWCS) that integrates literature-derived causal evidence with biological network context. At the corpus level, we introduce CWCS-TD, a truth discovery–based method specifically designed to operate under sparse biomedical evidence. CWCS-TD jointly estimates causal scores for G-D pairs and reliability scores for individual publications using bibliometric features. In parallel, we leverage curated gene regulatory networks and apply the CWCS-Net algorithm to capture network-driven causal influence that cannot be inferred from text alone.

By aggregating reliability-aware corpus-level evidence from CWCS-TD with network-based causal signals from CWCS-Net, CWCS (comb) achieves robust and consistent performance across diseases. Evaluation using OMIM-based validation demonstrates that CWCS attains an causal F1 score of 0.600 and PR-AUC of 0.601, outperforming standalone network-based, corpus-based, and large language model–based approaches. Although CWCS-TD alone does not exceed the performance of network-driven inference, its integration with CWCS-Net yields an absolute improvement of ~17% in PR-AUC, highlighting the complementary value of network based evidence.

While LLMs exhibit high recall in identifying potential causal mentions, they suffer from low precision when applied directly to corpus-level inference. In contrast, CWCS provides a stable framework that explicitly aggregates distributed evidence and accounts for source reliability, enabling more reliable causal gene prioritization under sparse evidence conditions.

Overall, this study demonstrates that integrating large-scale literature mining with biological network analysis offers a principled and effective approach for corpus-wide causal inference of G-D pairs. Future work will incorporate additional bibliometric and contextual features, and extend the framework to full-text articles with the goal of supporting causal discovery and target prioritization in precision medicine.

## Acknowledgments

We thank members of our BIRDS research group for their valuable inputs during the course of this work. We also thank Saish Jaiswal for his inputs on this manuscript. This work was supported by the Wellcome Trust/DBT India Alliance Intermediate Fellowship Grant IA/I/17/2/503323 awarded to MN. We would also like to acknowledge the use of LLMs for copy-editing certain sentences of the manuscript, and for obtaining code for certain minor helper functions, which were subsequently manually verified.

## 5. Appendix

### 5.1. Methods

#### 5.1.1. CWCS-TD Algorithm

##### T-step

The T-step is responsible for estimating the values 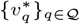, conditioned on the current estimates of source reliabilities.

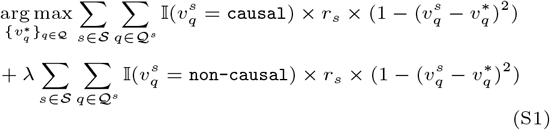

Equivalently, the objective can be written as,

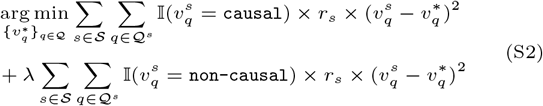

In effect, we are trying to optimize weighted distance of 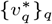, and so, it can be minimized by setting 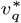 values to the weighted averages.

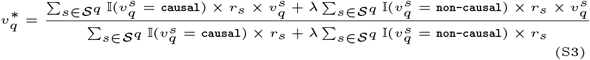

##### R-step

The R-step is responsible for estimating the reliability scores of sources, assuming access to the identified truths (as per Equation S3). As outlined in Section Extension of truth discovery algorithm (in the main text), source reliability is modeled as a linear combination of feature values. Consequently, estimating the weight vector {*β*_*f*_}_*f*_, which determines the contribution of each feature, is a prerequisite to computing the reliability scores.

Upon rearrangement, the objective (Equation S1) can be equivalently expressed as:

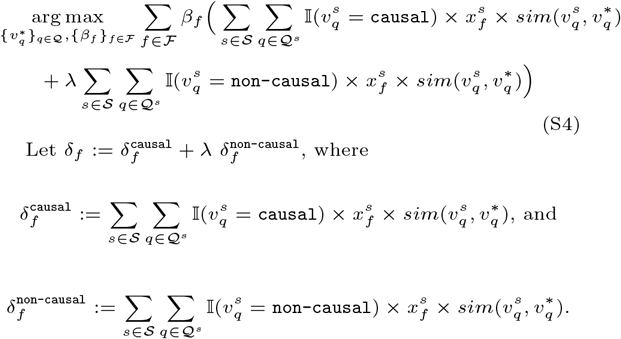

Therefore, the objective function in Equation S4 reduces to:

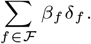

This rearrangement reveals an insight critical in defining the feasible region Δ when estimating *β* conditioned on the knowledge of the identified truths in the *R*−step. Evidently, the objective function is a linear combination of the weights {*β*_*f*_}_*f*_, and so constraining the feasible set such that Σ_*f*_ *β*_*f*_ = 1 with *β*_*f*_ ∈ [0, 1] ∀ *f* ∈ ℱ can lead to degenerate solutions. Specifically, the optimal solution would reduce to a one-hot vector—assigning *β*_*f*_ = 1 to the feature *f* ∈ ℱ that maximizes δ_*f*_, and zero to all others. In order to deal with such meaningless optimization, we define the feasible region for the reliability values as follows:

###### Algorithm 1

Source Aggregation and Reliability Estimation

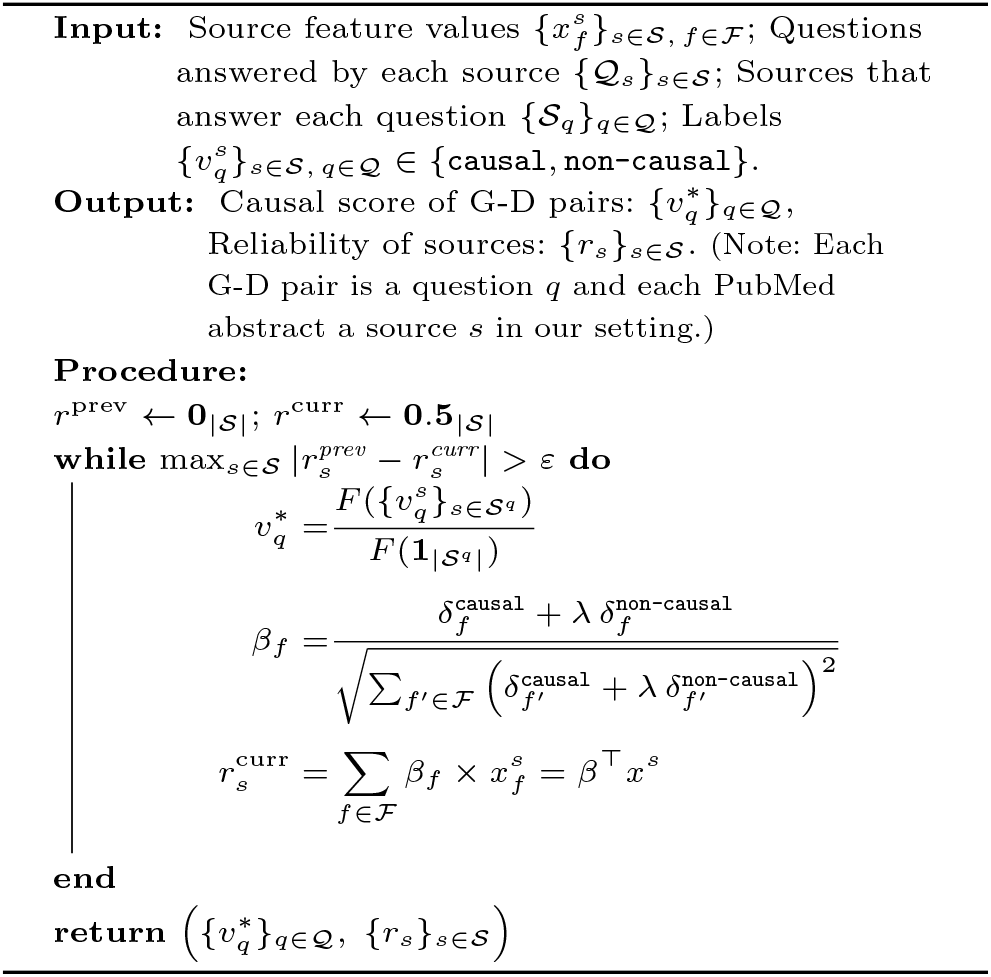

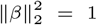

We now optimize for *β*, formally given as:

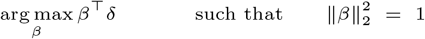

Geometrically, this optimization corresponds to maximizing the magnitude of the projection of the vector δ = {δ_*f*_}_*f*∈ℱ_ onto *β*, subject to the constraint that *β* lies on a unit hypersphere. The optimal solution is therefore achieved when *β* is perfectly aligned with *δ*, and is given by:

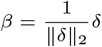

The notations used throughout the work are summarized in Table S1.

##### Algorithm Pseudocode

The pseudocode describing all the steps of the CWCS-TD algorithm is shown in Algorithm 1.

#### 5.1.2. LLM Inference: Prompt to LLM

Following prompt was provided to LLM for prediction: You are a biomedical expert. Your task is to determine whether a Causal relationship exists between @GeneSrc$ and @DiseaseTgt$ in overall text data provided to you. You will be provided with abstracts from various published papers and predictions from BioBERT+SVM for each abstract, including a label (1=Causal, 0=Not Causal) and a confidence score.

Instructions:

1. Analyze all the text evidence for explicit causal links.
2. Use the provided ‘Prior Model Predictions’ as supplementary signals.
3. Associations (e.g., “associated with”, “linked to”) are NOT causal. Look for explicit mention of causal relations like “causes”, “mutations in X lead to Y”, etc.
4. score is likelihood of causal relationship, its value depends on strength of evidence, try to be variable in score based on evidence.

Respond ONLY with valid JSON in this exact format:

“relationship”: “Causal” or “Not causal”,

“score”: float between 0 and 1,

“reasoning”: “Brief explanation”.

### 5.2. Supplementary Tables

**Table S1.**
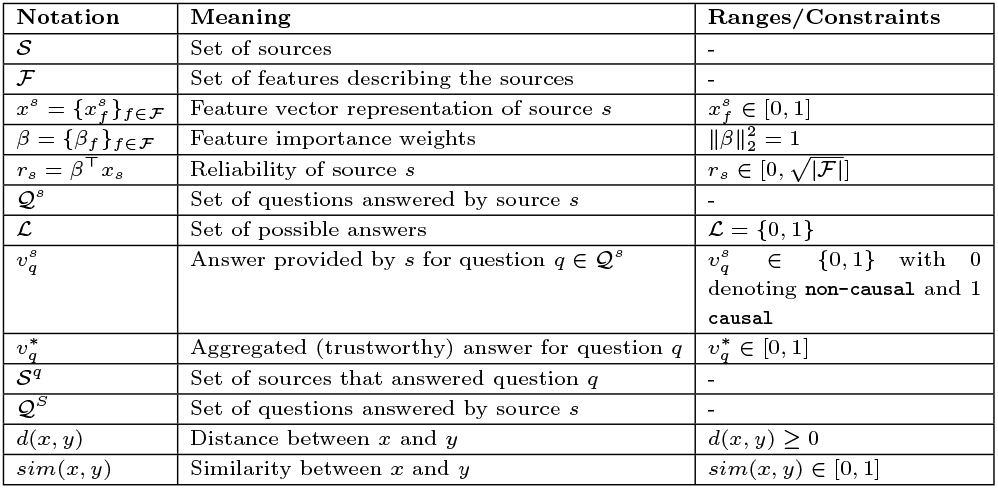
List of notations used, their meanings, and associated constraints.

**Table S2.** For each disease, the number of nodes and directed edges within each layer of its multilayer network is shown here.

| Disease | Nodes | G-D Edges | G-G Edges |
| --- | --- | --- | --- |
| Breast cancer | 260 | 259 | 2171 |
| Colorectal cancer | 111 | 110 | 841 |
| Gastric cancer | 22 | 21 | 46 |
| Hepatocellular carcinoma | 18 | 17 | 42 |
| Leukemia acute myeloid | 43 | 42 | 54 |
| Myocardial infarction | 36 | 35 | 32 |
| Ovarian cancer | 21 | 20 | 64 |
| Tetralogy of Fallot | 15 | 14 | 5 |
| Alzheimer Disease | 145 | 144 | 363 |
| Parkinson Disease | 203 | 202 | 528 |

**Table S3.**
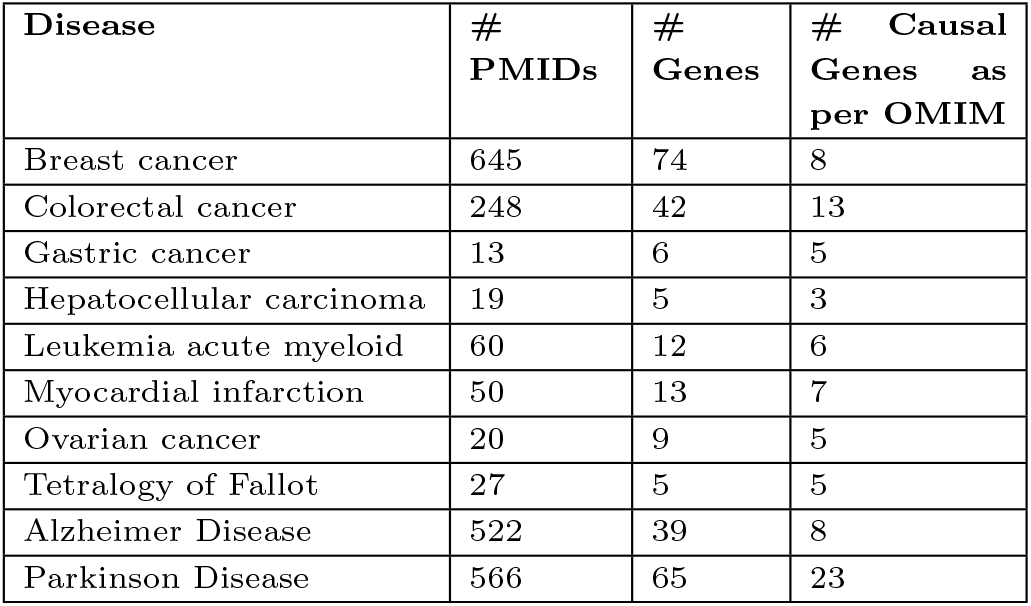
Statistics of the OMIM validation corpus when considering only those genes present in more than 1 paper as compared to Table 1 in the main text. # stands for “Number of”.

**Table S4.** Comparison of confusion matrix of all models.

| Model | TP | FP | FN | TN |
| --- | --- | --- | --- | --- |
| CWCS-TD | 65 | 102 | 18 | 84 |
| CausalRatio | 65 | 102 | 18 | 84 |
| GPT-4o | 78 | 148 | 5 | 38 |
| MMedLlama-3 | 60 | 87 | 23 | 99 |
| CWCS-Net | 62 | 84 | 21 | 102 |
| CWCS-Net+CausalRatio | 65 | 100 | 18 | 86 |
| CWCS-Net+GPT-4o | 59 | 81 | 24 | 105 |
| CWCS-Net+MMedLlama-3 | 67 | 102 | 16 | 84 |
| CWCS-Net+CWCS-TD | 54 | 43 | 29 | 143 |

**Table S5.**
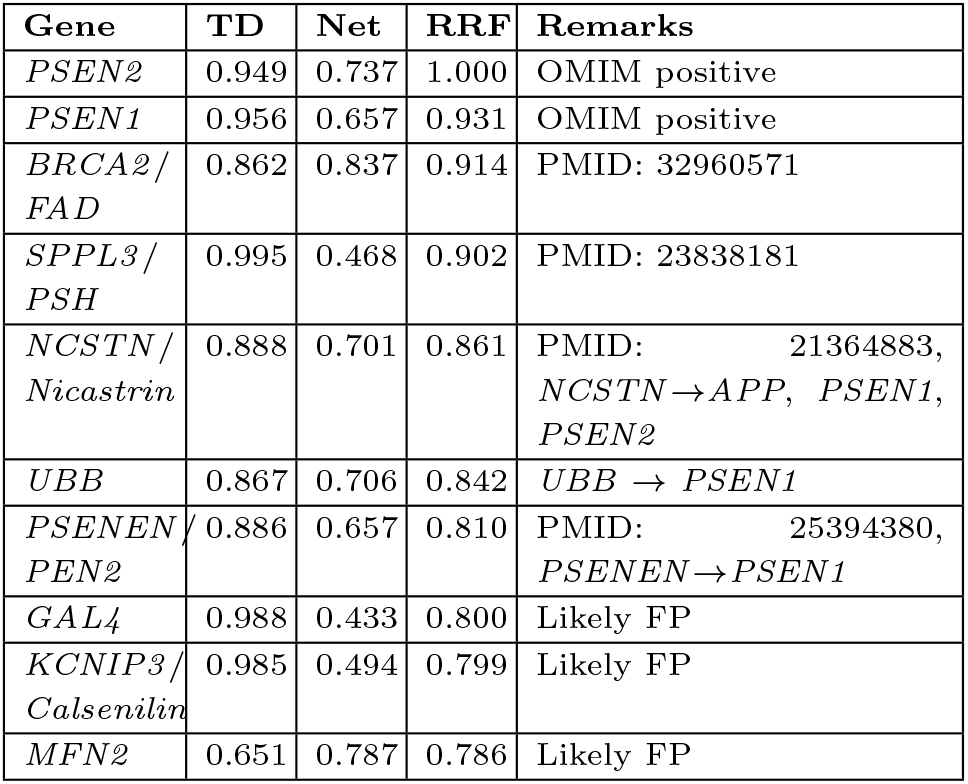
Top 10 genes with the highest CWCS (comb) scores from the full PubMed corpus for AD. They could be either captured in OMIM as a known causal gene for AD (marked as OMIM positive in the Remarks column), or could be potentially novel causal links (supported by textual evidence (PMID) or regulatory network edges, as indicated in the Remarks column), or likely false positives (FPs). TD, Net and RRF denote CWCS-TD, CWCS-Net and CWCS (comb) scores respectively.

| Gene | TD | Net | RRF | Remarks |
| --- | --- | --- | --- | --- |
| *PSEN2* | 0.949 | 0.737 | 1.000 | OMIM positive |
| *PSEN1* | 0.956 | 0.657 | 0.931 | OMIM positive |
| *BRCA2/*<br>*FAD* | 0.862 | 0.837 | 0.914 | PMID: 32960571 |
| *SPPL3/*<br>*PSH* | 0.995 | 0.468 | 0.902 | PMID: 23838181 |
| *NCSTN/*<br>*Nicastrin* | 0.888 | 0.701 | 0.861 | PMID: 21364883,<br>*NCSTN*→*APP*, *PSEN1*,<br>*PSEN2* |
| *UBB* | 0.867 | 0.706 | 0.842 | *UBB* → *PSEN1* |
| *PSENEN/*<br>*PEN2* | 0.886 | 0.657 | 0.810 | PMID: 25394380,<br>*PSENEN*→*PSEN1* |
| *GAL4* | 0.988 | 0.433 | 0.800 | Likely FP |
| *KCNIP3/*<br>*Calsenilin* | 0.985 | 0.494 | 0.799 | Likely FP |
| *MFN2* | 0.651 | 0.787 | 0.786 | Likely FP |

**Table S6.** Top 10 genes with the highest CWCS (comb) scores from the full PubMed corpus for PD. They could be either captured in OMIM as a known causal gene for PD (marked as OMIM positive in the Remarks column), or could be potentially novel causal links (supported by textual evidence (PMID) or regulatory network edges, as indicated in the Remarks column), or likely false positives (FPs). TD, Net and RRF denote CWCS-TD, CWCS-Net and CWCS (comb) scores respectively.

| Gene | TD | Net | RRF | Remarks |
| --- | --- | --- | --- | --- |
| *RAB32* | 0.872 | 1.000 | 1.000 | OMIM positive |
| *DNAJC6* | 0.945 | 0.801 | 0.987 | OMIM positive |
| *GBA1* | 0.894 | 0.795 | 0.946 | OMIM positive |
| *FBXO7* | 0.867 | 0.766 | 0.903 | OMIM positive |
| *NR4A2* | 0.744 | 0.876 | 0.894 | OMIM positive |
| *UCHL1* | 0.649 | 0.906 | 0.879 | OMIM positive |
| *EIF4G1* | 0.849 | 0.765 | 0.876 | OMIM positive |
| *HTRA2* | 0.811 | 0.768 | 0.861 | OMIM positive |
| *PARK7* | 0.698 | 0.806 | 0.856 | OMIM positive |
| *IL10RA* | 0.997 | 0.391 | 0.852 | Likely FP |

**Table S7.** Manually verified examples of PMIDs. These PMIDs are the ones with high reliability score for the OMIM-listed causal genes of AD. Label (causal/ non-causal) is assigned by the manual annotator. For some genes, the mentioned snippets of abstracts might not be sufficient to decide the label; so we considered the complete abstract while annotating it.

| Gene | PMID | Label | Snippets from abstracts |
| --- | --- | --- | --- |
| *APP* | 1302033 | causal | "Mutations at codon 717 in exon 17 of the beta-amyloid precursor protein (APP) gene have previously been shown to segregate with early onset Alzheimer's disease in some families." |
| *APP* | 1280524 | non-causal | "The presence of beta APP in dystrophic neurites of SP and the localization of beta APP in lysosomes suggest that beta APP containing dystrophic neurites may play a role in the extracellular deposition of amyloid." |
| *APP* | 1303239 | causal | "Several families with an early-onset form of familial Alzheimer's disease have been found to harbour mutations at a specific codon (717) of the gene for the beta-amyloid precursor protein (APP) on chromosome 21." |
| *APOE* | 8446617 | causal | "Analysis of apolipoprotein E alleles in Alzheimer disease and controls demonstrated that there was a highly significant association of apolipoprotein E type 4 allele (APOE-epsilon 4) and late-onset familial Alzheimer disease." |
| *APOE* | 7972031 | causal | "ApoE is found in some neurofibrillary tangle-bearing neurons, one of the major pathologic hallmarks of the disease." |
| *APOE* | 12198535 | non-causal | "This effect of the APOE epsilon 4 allele on normal cognitive ageing may be mediated by a mechanism that is at least partly independent of its predisposing effect towards Alzheimer's disease." |
| *PSEN1* | 8799182 | causal | "Mutations in the recently identified presenilin 1 gene on chromosome 14 cause early onset familial Alzheimer disease (FAD)." |
| *PSEN1* | 8943053 | causal | "Presenilin-1 (PS-1) gene mutations are responsible for the majority of the early onset familial forms of Alzheimer disease (AD)." |
| *PSEN1* | 8962160 | non-causal | "The reduced activity of mutant presenilins and as yet unknown gain-of-function properties may be a contributing factor in the development of Alzheimer disease." |

**Table S8.**
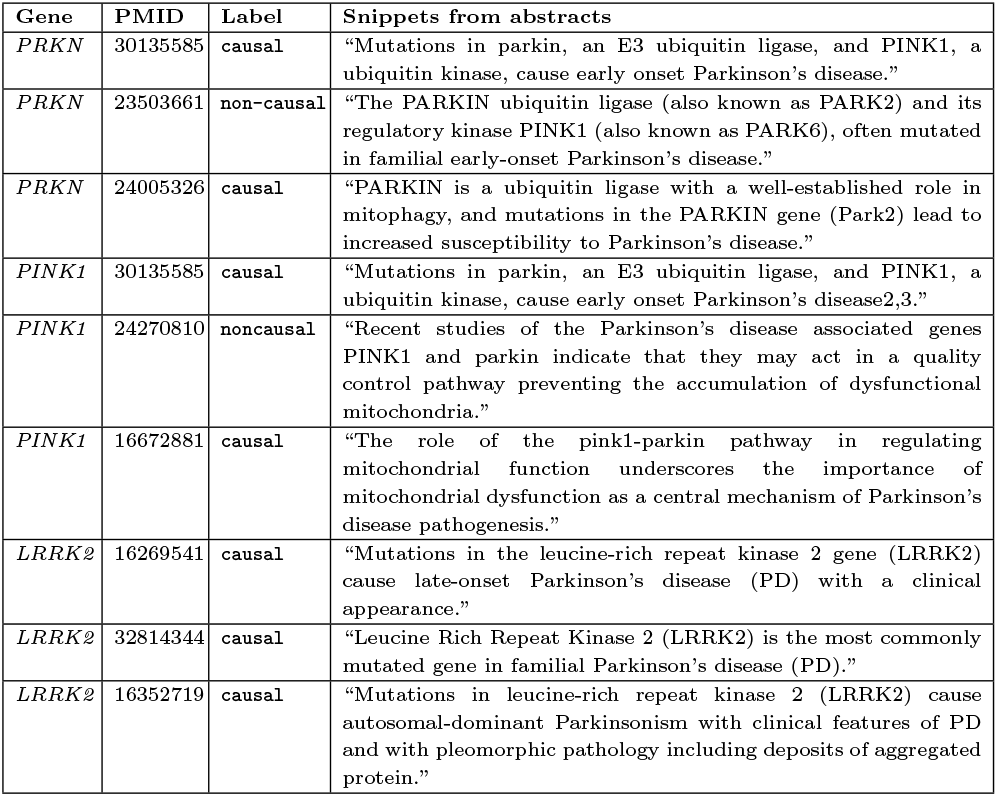
Manually verified examples of PMIDs. These PMIDs are the ones with high reliability score for the OMIM-listed causal genes of PD. Label (causal/ non-causal) is assigned by the manual annotator. For some genes, the mentioned snippets of abstracts might not be sufficient to decide the label; so we considered the complete abstract while annotating it.

### 5.3. Supplementary Figures

**Fig. S1.**
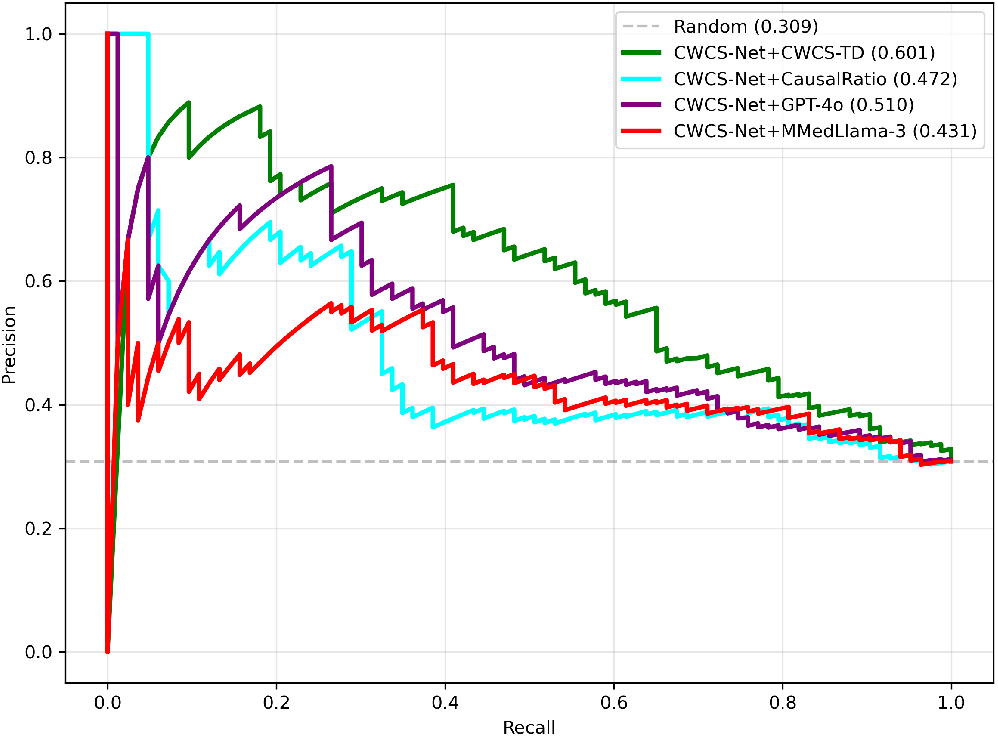
PR curve of different models aggregated with CWCS-Net scores on OMIM validation corpus.

**Fig. S2.**
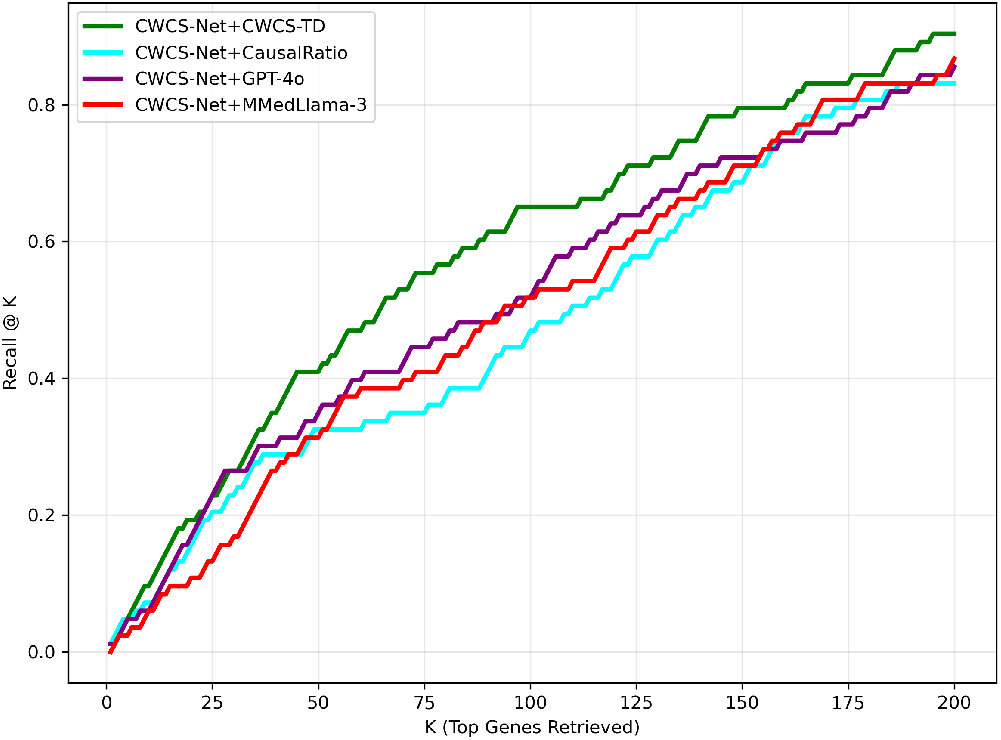
Recall@K curve of different models aggregated with CWCS-Net scores on OMIM validation corpus.

